# *XIST* Is a Key Modulator Associated with the Adhesome Network

**DOI:** 10.64898/2026.04.21.719966

**Authors:** Dongning Chen, Nolan Origer, Sha Sun, Timothy L. Downing

**Affiliations:** Department of Developmental and Cell Biology, University of California, Irvine, CA 92697, USA; Department of Biomedical Engineering, University of California, Irvine, CA 92697, USA; Department of Systems Biology, University of California, Irvine, CA 92697, USA; Department of Microbiology and Molecular Genetics, University of California, Irvine, CA 92697, USA

**Keywords:** lncRNA, *XIST*, adhesome, stemness, epigenetics

## Abstract

A long non-coding RNA (lncRNA) known as the X-inactivation specific transcript (*XIST*) plays a central role in X chromosome inactivation – a transcriptional process that silences one of the two X chromosomes in females to ensure dosage compensation between males and females. Much research has been conducted on how *XIST* regulates X chromosome transcription critical to embryonic development, but recent studies suggest a non-canonical role for *XIST* in regulating cancer stem cells and cellular plasticity. As cell adhesion and adhesome genes are integral to the regulation of cancer stemness, we explored the previously unrecognized link between *XIST* and the adhesome network. By performing gene expression and gene ontology analysis on *XIST*-knockdown ovarian cancer cells, our study showed that *XIST* loss altered adhesome gene expression and downstream adhesion pathways. Using Genotype-Tissue Expression (GTEx) and The Cancer Genome Atlas (TCGA) datasets, we identified distinct correlations between *XIST* lncRNA and adhesome genes across normal and cancer tissue samples, which are associated with cell stemness. Furthermore, network analysis suggests that *XIST* may interact with specific adhesome genes within the cell nucleus. This interaction may have significant functional implications, as demonstrated by the hazard ratio analysis of *XIST* and adhesome gene expression in relation to clinical outcomes. Overall, our results show that among well-annotated functional lncRNAs, *XIST* appears to be a modulator strongly associated with the adhesome network and cell stemness. Our findings thus support a novel link between lncRNA-mediated epigenetic regulation of cell adhesion genes, highlighting *XIST* as a key regulator contributing to the adhesome network.

**Significance Statement:** This study identified that *XIST*, a long non-coding RNA essential for X-chromosome dosage compensation and embryonic development, plays a significant role in modulating the adhesome network. We found that *XIST* knockdown affected adhesion pathways in ovarian cancer cells, whereas *XIST* expression is strongly correlated with adhesome gene expression across all tissues. We observed that the interaction between *XIST* and adhesome genes changes significantly between tumors and normal tissues, and this altered interaction is associated with certain cancer outcomes. These findings reveal a possible link between lncRNA-mediated regulation and adhesome control that is associated with cell stemness signatures and the emergence of cancerous tissues.

## Introduction

Non-coding RNAs (ncRNAs; RNAs that do not encode proteins) play critical roles in gene regulation and are central to epigenetics (1–4). Long non-coding RNAs (lncRNAs) are larger than 200 nucleotides and are important building blocks of gene regulatory networks in all eukaryotes (5–7). Mammalian lncRNAs have been shown to affect early development and certain diseases like cancer can be caused by dysregulation of human lncRNAs; however, much is still unknown about how lncRNAs regulate cellular physiology and disease (8–11). The X-inactivation specific transcript (*XIST*) is a 17-kb lncRNA critical for embryonic development while eliciting transcriptional inactivation of one of the two X chromosomes that is maintained in all somatic cells throughout a female’s lifespan (12, 13). Aside from its established function in X-chromosome inactivation, *XIST* dysregulation has been implicated in a variety of cancers, where it plays a non-canonical role in tumorigenesis and progression (14–16). Importantly, *XIST* loss disrupts cell differentiation and enhances stemness (16, 17), and recent evidence suggests that *XIST* regulates autosomal gene expression in pluripotent cells, extending its transcriptional control beyond the X chromosome (18).

Our previous work identified that *XIST* knockdown in human ovarian cancer OVCAR3 cells results in cellular changes that affect cell adhesion, migration, invasion and epithelial-mesenchymal transition (EMT) (16). These findings suggested a possible link between *XIST* and the cell adhesion process, motivating further investigation into its molecular mechanisms. Cell adhesion is regulated by the adhesome, characterized as a network of hundreds of genes encoding proteins that mediate cell-cell and cell-extracellular matrix (ECM) interactions, cytoskeletal organization, and intercellular signaling (19, 20). Adhesome proteins, including their immediate downstream signaling partners, facilitate two major types of adhesion: cell-matrix adhesion, mediated by integrins and focal adhesion kinases, which anchor cells to the extracellular matrix; and cell-cell adhesion, mediated by cadherins and catenins, which maintain intercellular contact and tissue integrity (21, 22). Catenins, including β-catenin, function as adaptor proteins that link the intracellular domain of cadherins to actin, stabilizing adherens junctions (23). When incorporated into cadherin-β-catenin-actin complexes, β-catenin is sequestered from nuclear Wnt signaling, whereas Wnt pathway activation redirects β-catenin from adhesion complexes to the nucleus, thereby altering adhesion dynamics (24, 25). As with *XIST*, adhesion and other biophysical interactions contribute to stemness acquisition and gene expression regulation in pluripotent cells (26–29). The adhesome plays a central role in guiding cell fate decisions and behaviors (19, 30–32), and it has also been implicated in more malicious cell fates, such as tumorigenesis (33–37). Many processes within the tumor microenvironment, such as growth, angiogenesis, and metastasis, rely on communication and signaling between cells via physical interactions with each other and their environment, and can be significantly affected by gain- or loss-of-function mutations in the adhesome, which are common in cancer pathology (36, 38, 39). Dysregulation of the adhesome in cancer disrupts cell adhesion, enhances focal adhesion turnover, and promotes cell motility, which further drives epithelial-mesenchymal transition (EMT) and tumor invasion (40, 41). Moreover, adhesome dysregulation is closely linked to cancer stemness, although the contribution of biophysical cues varies across cellular contexts (42–44).

Based on these observations, we hypothesized that *XIST* may influence the adhesome network, revealing a novel link between lncRNA-mediated epigenetic regulation and cell adhesion pathways. To test this hypothesis, we investigated transcriptional changes in adhesome genes following *XIST* knockdown, evaluated correlations between the expression of lncRNAs and adhesome genes, and analyzed *XIST*-adhesome correlations with cell stemness. We further constructed an *XIST*-associated adhesome network using potentially nuclear-localized adhesome genes, which showed increased correlation with *XIST* expression. We found that a subset of adhesome genes and multiple adhesion-related pathways were altered following the loss of *XIST*. Expression correlation analysis positioned *XIST* as the top lncRNA potentially regulating adhesome genes with strong correlations across normal tissues that diminished in tumor samples. Furthermore, the interaction between *XIST* and adhesome genes is strongly associated with cell stemness and affects cancer outcomes. By identifying the lncRNA *XIST* as a potential key modulator associated with the adhesome network, our study establishes a new framework for understanding how cell adhesion and epigenetic mechanisms interact to regulate cell fate and tumor progression.

## Results

### *XIST* knockdown alters adhesome expression and adhesion-related pathways

To investigate the potential link between *XIST* and adhesome genes, we analyzed differential gene expression in human ovarian cancer cells (OVCAR3) following *XIST* knockdown under normoxic conditions. Two independent knockdown clones, X7 (sgXIST7) and X9 (sgXIST9), were generated using distinct guide RNAs targeting different regions of the *XIST* promoter (sgXIST7: 5’-GCAGCGCTTTAAGAACTGAA-3’; sgXIST9: 5’-GCCATATTTCTTACTCTCTCG-3’), thereby minimizing potential guide-specific off-target effects (16). Knockdown efficiency was validated by examining *XIST* RNA-seq read counts across all samples (Supplementary Table 3). Control samples showed robust *XIST* expression (4,900, 11,209, and 4,213 reads), whereas *XIST* expression was nearly abolished in the X7clone (5, 4, and 1 reads) and substantially reduced in the X9 clone (650, 1,009, and 898 reads), consistent differing knockdown efficiencies of the two guide RNAs. To assess overall transcriptomic effects, we performed principal component analysis (PCA) on all samples. Control samples separated clearly from both X7 and X9 clones along PC1 (69% variance), indicating global transcriptional changes associated with *XIST* depletion (Supplementary Figure 1e). X7 and X9 clones further separated along PC2 (23% variance), consistent with their differential knockdown efficiencies. Building on this initial assessment, we analyzed the previously generated DESeq2 dataset (16), which included X7 (n = 3), X9 (n = 3), and control samples (n = 3) under normoxic conditions, to identify differentially expressed genes (DEGs) (Figure 1a). DEGs were defined using a fold-change threshold of 1.5 (|log2_FoldChange| > 0.58) and adjusted p value < 0.05 (Supplementary Table 1). In the X7 clone, 2,509 DEGs were identified, including 68 adhesome genes – representing a significant enrichment relative to the 334 curated adhesome genes (19, 20) (p = 0.0062, hypergeometric test; Figure 1b, left panel; Supplementary Figure 1a). Similarly, in the X9 clone, 64 of 2,563 DEGs were adhesome genes, also showing significant enrichment (p = 0.0401, hypergeometric test; Figure 1b, middle panel; Supplementary Figure 1b). Together, these results demonstrate a significant association between *XIST* depletion and altered adhesome gene expression.

**Figure 1.**
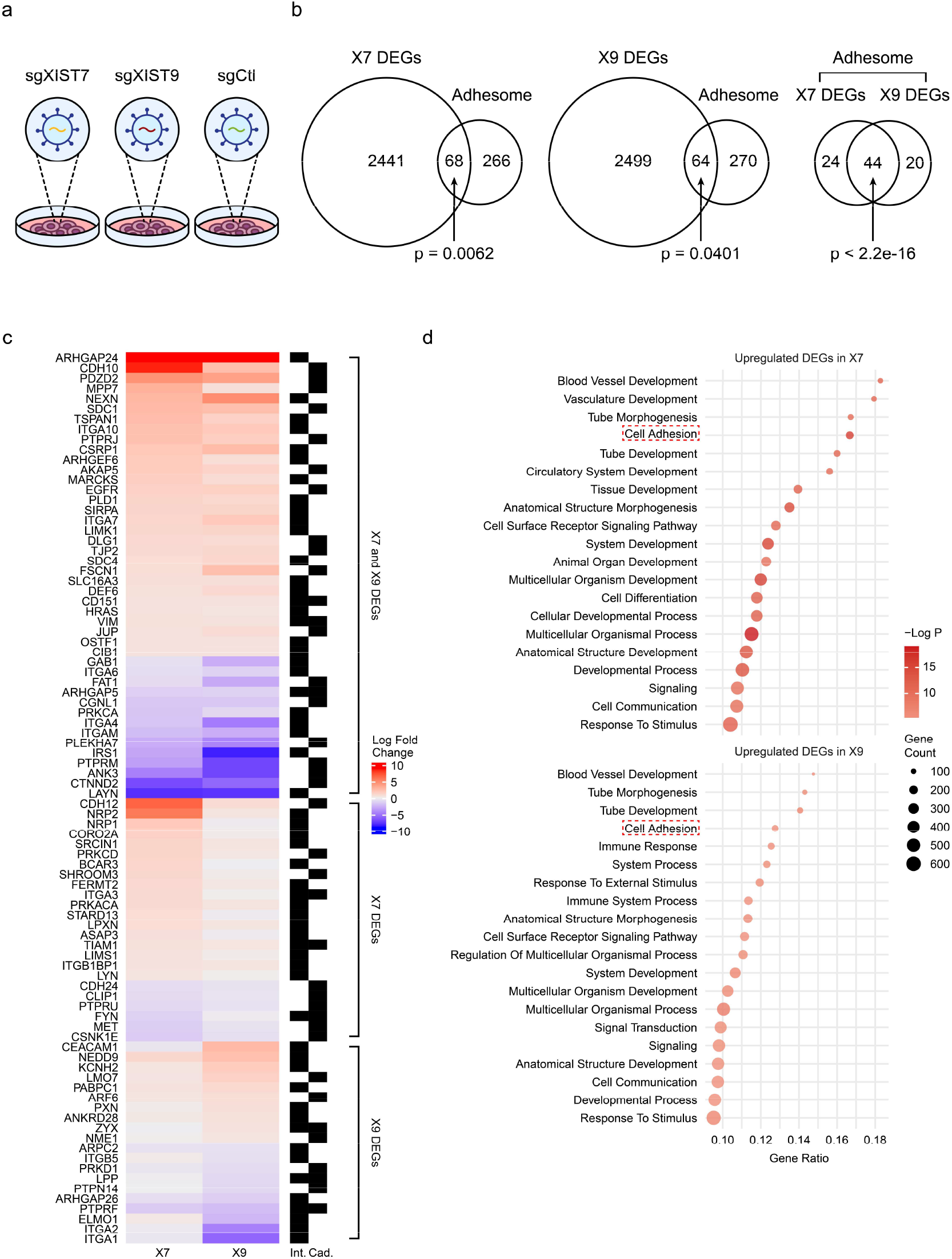
Adhesome gene expressions and adhesion-related biological processes are significantly altered following *XIST* loss in OVCAR3. (a) OVCAR3 *XIST* knockdown clones X7 (n = 3) and X9 (n = 3), and OVCAR3 control clones (n = 3). (b) Venn diagram illustrating the significant overlap between adhesome genes and DEGs derived from *XIST* knockdown OVCAR3 clone X7 (left panel) or X9 (middle panel), and the significant overlap between adhesome genes differentially expressed in OVCAR3 clone X7 and X9 (right panel). Statistical analysis was performed by a hypergeometric test. (c) A heatmap of differentially expressed adhesome genes following *XIST* knockdown and their classification (integrin vs cadherin associated). The color represents log_2_ fold change values, with upregulated genes shown in red and downregulated genes in blue. (d) The top 20 enriched GO Biological Process (GOBP) terms based on upregulated DEGs in the X7 and X9 clone. GO analysis was performed using the gprofiler2 R package. Each dot size represents the count of genes associated with the term and the color indicates the statistical significance (-log_10_(p value)). The x-axis represents the gene ratio, defined as the proportion of genes enriched in each term relative to the total number of genes in that gene set. Terms were presented by ranking by their gene ratios.

Across both X7 and X9 clones, 44 adhesome genes were consistently differentially expressed (Figure 1b, right panel), all showing concordant transcriptional changes following *XIST* loss (Figure 1c). Notably, adhesome genes such as *ARHGAP24, CDH10*, and *PDZD2* were upregulated in both clones, whereas *LAYN, CTNND2*, and *ANK3* were downregulated (Figure 1c). Upregulated adhesome genes were significantly enriched in both X7 (p = 0.00045) and X9 (p = 0.02997), indicating activation of adhesome-related transcriptional programs in response to *XIST* depletion (Supplementary Figure 1c-d). Because upregulated adhesome genes substantially outnumbered downregulated adhesome genes in both knockdown clones (X7: 48 upregulated vs. 20 downregulated; X9: 40 upregulated vs. 24 downregulated) and showed greater consistency between X7 and X9, we focused subsequent pathway enrichment analyses on the upregulated DEGs. Gene ontology enrichment analysis showed that upregulated DEGs in both knockdown clones were significantly enriched for the cell adhesion pathway (Figure 1d).

Additional enriched terms included several developmental and morphogenesis-related processes that are closely linked to adhesome function, such as epithelial organization and tissue remodeling (Figure 1d). These processes are driven by genes involved in cell-cell junctions, extracellular matrix organization, migration, and polarity – core components of the adhesome network – and therefore likely reflect broader changes in adhesion-related architecture following *XIST* loss. Whereas the prior study emphasized stemness- and EMT-associated programs, the present work specifically centers on the adhesome network. Our results indicate that *XIST* loss influences adhesome gene expression and modulates adhesion-related pathways.

### *XIST* strongly correlates with adhesome expression in normal tissues but not in tumors

*XIST* is a female-specific lncRNA that is expressed in all female somatic tissues. To assess the dynamic relationship between *XIST* and adhesome genes, we performed expression correlation analysis using female samples from the Genotype-Tissue Expression (GTEx) and The Cancer Genome Atlas (TCGA) datasets (45, 46). The expression levels of *XIST* across tissue types in these databases are shown in Supplementary Figure 2a-b, providing context for the correlation analyses. Among functional lncRNAs with well-defined annotations and reported roles in gene regulation, *XIST* exhibited the strongest correlation with adhesome genes across normal tissues – significantly exceeding that of other representative lncRNAs analyzed, including *GAS5* and *MALAT1*, which may be involved in adhesome-related pathways (47–49) (Figure 2a, green curves; Supplementary Figure 3a). The comparator lncRNAs were selected as a set of well-characterized regulatory lncRNAs with diverse reported functions, providing a biologically relevant reference framework for evaluating relative correlation strength rather than implying shared mechanistic roles with *XIST*. Within this lncRNA panel, *XIST* showed a distinct and consistently stronger association with adhesome gene expression, identifying it as a particularly strong candidate regulator within the adhesome network.

**Figure 2.**
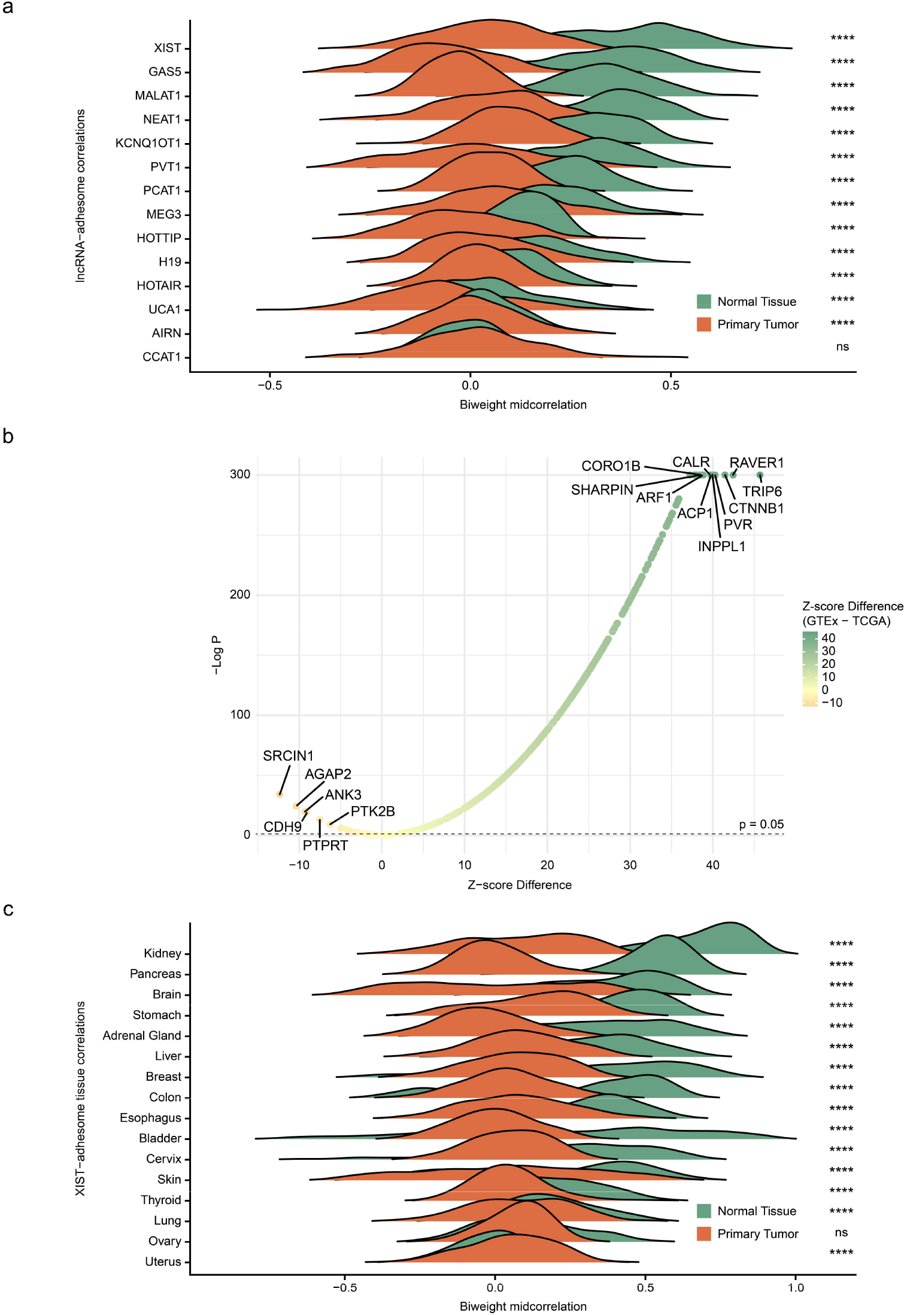
*XIST* strongly correlates with adhesome genes in normal tissues, with reduced correlations in primary tumors. (a) The correlation (calculated by biweight midcorrelation) between lncRNAs and adhesome genes in normal tissues (female, from GTEx) and primary tumors (female, from TCGA). Statistical analysis was performed by one tailed Wilcoxon test. (b) The differential correlation of *XIST*-adhesome expression between normal and tumor states. (c) The comparison of correlation between *XIST* and the adhesome in shared tissue types in normal tissues and in primary tumors. ns, not significant; ****, p < 0.0001.

To assess whether the observed *XIST*-adhesome correlation could occur by chance, we performed a permutation analysis using 2,000 random gene sets of the same size as the adhesome set, sampled either from the entire genome or restricted to X-linked genes. The median correlation between *XIST* and adhesome genes (R = 0.3993) exceeded that of all random sets, demonstrating that the strong positive association is highly specific (Supplementary Figure 2c-d). Supporting this finding in an independent system, our analysis of mouse tissue expression profiles from the Gene Expression Database (GXD) with Mouse Genome Informatics (MGI) (50), showed that the mouse lncRNA *Xist* also displays strong correlations with adhesome genes (Supplementary Figure 3e). Although the correlation strength of *Xist* was comparable to that of *Kcnq1ot1* and *Gas5*, it remained significantly higher than that of other lncRNAs examined (Supplementary Figure 3e), suggesting a conserved role for *XIST/Xist* associated with adhesome regulation.

In contrast, the *XIST*-adhesome correlations were significantly reduced in primary tumors (Figure 2a; Supplementary Figure 3b). A differential comparison of *XIST*-adhesome correlations between normal versus tumor tissues showed that most adhesome genes had significantly higher correlations with *XIST* in normal tissues than in tumors (Figure 2b). Adhesome genes such as *TRIP6, RAVER1* and *CTNNB1* were highly correlated with *XIST* in normal tissues but lost their strong association in tumors. These shifts indicate that the relationship between *XIST* and adhesome gene expression is dynamic and influenced by disease state. When we examined individual tissue types with matched normal and tumor samples, we found that *XIST*-adhesome correlation strength varied across tissues (Supplementary Figure 3c-d). Normal tissues such as kidney, pancreas, heart, and brain showed robust positive correlations between *XIST* and adhesome, with most values falling in the moderate-to-high range (Supplementary Figure 3c). In contrast, most tumor tissues showed weaker or even negative correlations (Supplementary Figure 3d). Nonetheless, certain tumor tissues such as lymphatic tissue and thymus retained relatively strong positive correlations (Supplementary Figure 3d). Analysis of shared tissue types between GTEx and TCGA further confirmed that, for most tissues, *XIST*-adhesome correlations were significantly higher in normal states than in tumors (Figure 2c). The ovary showed no difference between states, whereas the uterus exhibited stronger correlations in tumors than in normal tissues. These findings demonstrate that the regulatory relationship between *XIST* and the adhesome is substantially altered in cancer and is highly tissue-dependent under both normal and tumor conditions.

In summary, expression correlation analysis suggests that *XIST* may be part of the adhesome network as a regulatory lncRNA, and that its interactions with adhesome genes are tissue-dependent and can be disrupted in cancer.

### *XIST* preferentially associates with nuclear adhesome genes and affects cancer prognosis

Adhesome components are traditionally defined as the network of proteins that connect the extracellular matrix and actin cytoskeleton at adhesion sites on the plasma membrane; however, research has identified that some adhesome proteins can also localize to the nucleus, with adhesome components shuttling between the cytoplasm and the nucleus in response to mechanical or signaling stimuli (51, 52). Using SubcellulaRVis, a bioinformatics tool that infers protein subcellular localization based on Gene Ontology Cellular Component (GOCC) enrichment analysis (53), we identified and classified adhesome genes as potentially nuclear or non-nuclear localized (Supplementary Table 2). We found that *XIST*, a nuclear-localized lncRNA, showed significantly stronger correlations with potentially nuclear-localized adhesome genes compared to non-nuclear localized ones, a pattern not observed in primary tumors (Figure 3a). This finding suggests that co-regulation of lncRNA *XIST* with the adhesome may rely on direct protein interactions localized to the nucleus, and that such regulatory interactions are altered during tumor progression. Based on these profiles, we constructed an *XIST*-associated adhesome protein-protein network using the top 10% of (potentially) nuclear adhesome genes ranked by correlation strength with *XIST. CTNNB1* (β-catenin) emerged as the dominant hub, with *ABL1* forming a secondary node within this network (Figure 3b). A regulatory interaction between *XIST* and β-catenin has been reported in the context of normal cellular physiology, supporting the biological relevance of this *CTNNB1*-centered network (54, 55).

**Figure 3.**
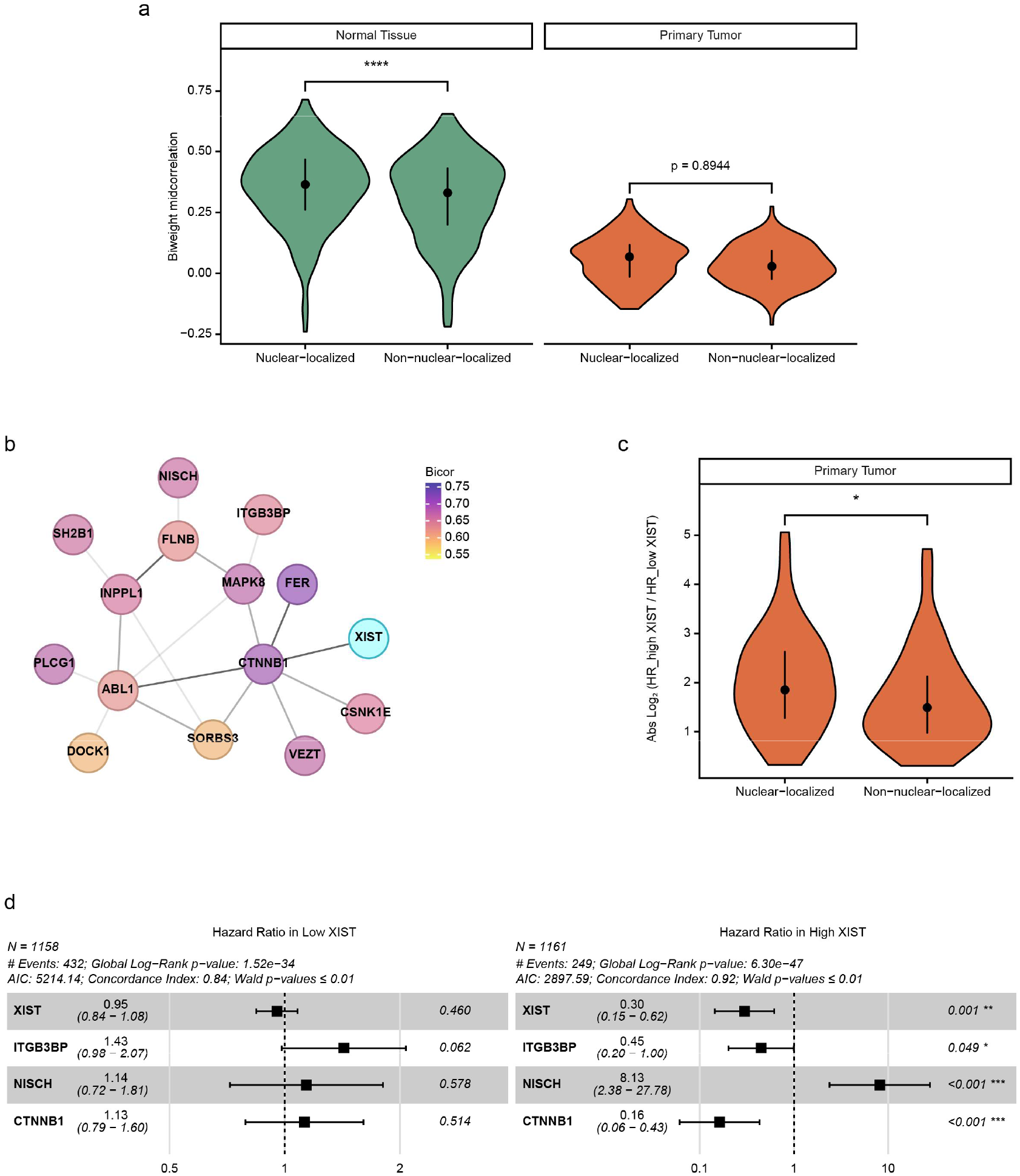
*XIST* is highly associated with an adhesome network, potentially through interaction with *CTNNB1*. (a) Violin plots comparing correlations between *XIST* and adhesome genes predicted to be potentially nuclear-localized versus non-nuclear-localized in normal tissue and primary tumor. Data normality was assessed using the Shapiro-Wilk test and statistical analysis was performed by Wilcoxon test (left panel) and unpaired t test (right panel). (b) *XIST*-associated adhesome network constructed using STRING and visualized in Cytoscape, based on nuclear adhesome genes with top 10% correlation with *XIST*. (c) Violin plots comparing the absolute log_2_ HR ratios (log_2_(HR_high *XIST* / HR_low *XIST*)) between high and low *XIST* samples in nuclear-localized and non-nuclear-localized adhesome genes. Data normality was assessed using the Shapiro-Wilk test and statistical analysis was performed by Wilcoxon test. (d) HR of network adhesome genes with significant HR changes (Wald p < 0.05) in low and high *XIST* pan-cancer samples. *, p < 0.05; **, p < 0.01; ***, p < 0.001; ****, p < 0.0001.

Using the TCGA primary tumor pan-cancer dataset with available patient survival data, we evaluated how *XIST* expression influences the relationship between adhesome gene expression and clinical outcomes. For each adhesome gene, a hazard ratio (HR) was calculated for tumors in the highest (High) and lowest (Low) quartiles of *XIST* expression levels. The effect of *XIST* expression on each adhesome gene was then quantified as the absolute log_2_ ratio of HRs between High-*XIST* and Low-*XIST* group (Supplementary Table 2). Among the 334 adhesome genes, 89 showed significant *XIST*-dependent changes in hazard ratio (Wald p < 0.05). The effect of *XIST* was stronger for nuclear-localized adhesome genes compared to non-nuclear-localized ones (Figure 3c), supporting a nuclear role for *XIST* within the adhesome network.

Consistent with these observations, *XIST* itself exhibited differential risk values between High-*XIST* and Low-*XIST* groups, with higher *XIST* activity associated with a more protective effect against tumor progression. Among the key candidates in the *XIST*-associated adhesome network, *CTNNB1, ITGB3BP*, and *NISCH* showed significant *XIST*-dependent changes in HR (Figure 3d). Both *CTNNB1* and *ITGB3BP* displayed reduced HR within the High-*XIST* cohort, indicating a potential protective role for these genes against worse cancer outcomes. *NISCH* exhibited the opposite trend, within the High-*XIST* context where its expression was associated with HR greater than 1, suggestive of pro-tumorigenic effect. These effects appeared specific to the High-*XIST* cohort as no significant changes in HR were observed within Low-*XIST* samples. Collectively, these results indicate that nuclear adhesome genes strongly influence cancer patient outcomes in a manner associated with *XIST* expression, with *XIST* potentially interacting with *CTNNB1* to strengthen tumor suppressive pathways.

Altogether, these findings suggest that *XIST*-adhesome interactions occur preferentially within the cell nucleus, particularly within in a *CTNNB1*-centered network, where they may help stabilize adhesion dynamics under physiological conditions, whereas disruption of these interactions in tumors adversely affects patient outcomes.

### *XIST* and adhesome expression are inversely correlated with stemness under both normal and tumor conditions

Given our previous finding that loss of *XIST* increases stemness in ovarian cancer cells (16), we next examined the relationship between *XIST* expression and stemness across normal, primary tumor, and metastatic states. *XIST* showed a negative correlation with stemness in all three conditions (Figure 4a). This correlation was strongest in normal tissues (R = -0.5583) and weakened in primary tumors (R = -0.1856) and metastatic samples (R = -0.2357), suggesting that the inhibitory relationship between *XIST* and stemness is attenuated in tumors. When samples were stratified into the highest (High) and lowest (Low) quartiles of *XIST* expression, High-*XIST* samples consistently exhibited significantly lower stemness scores across all three conditions (Figure 4b). These results indicate that *XIST* is consistently associated with the suppression of stemness, and that reduced *XIST* expression may promote the acquisition of cancer stem-like properties, potentially enhancing tumor aggressiveness and metastatic potential.

**Figure 4.**
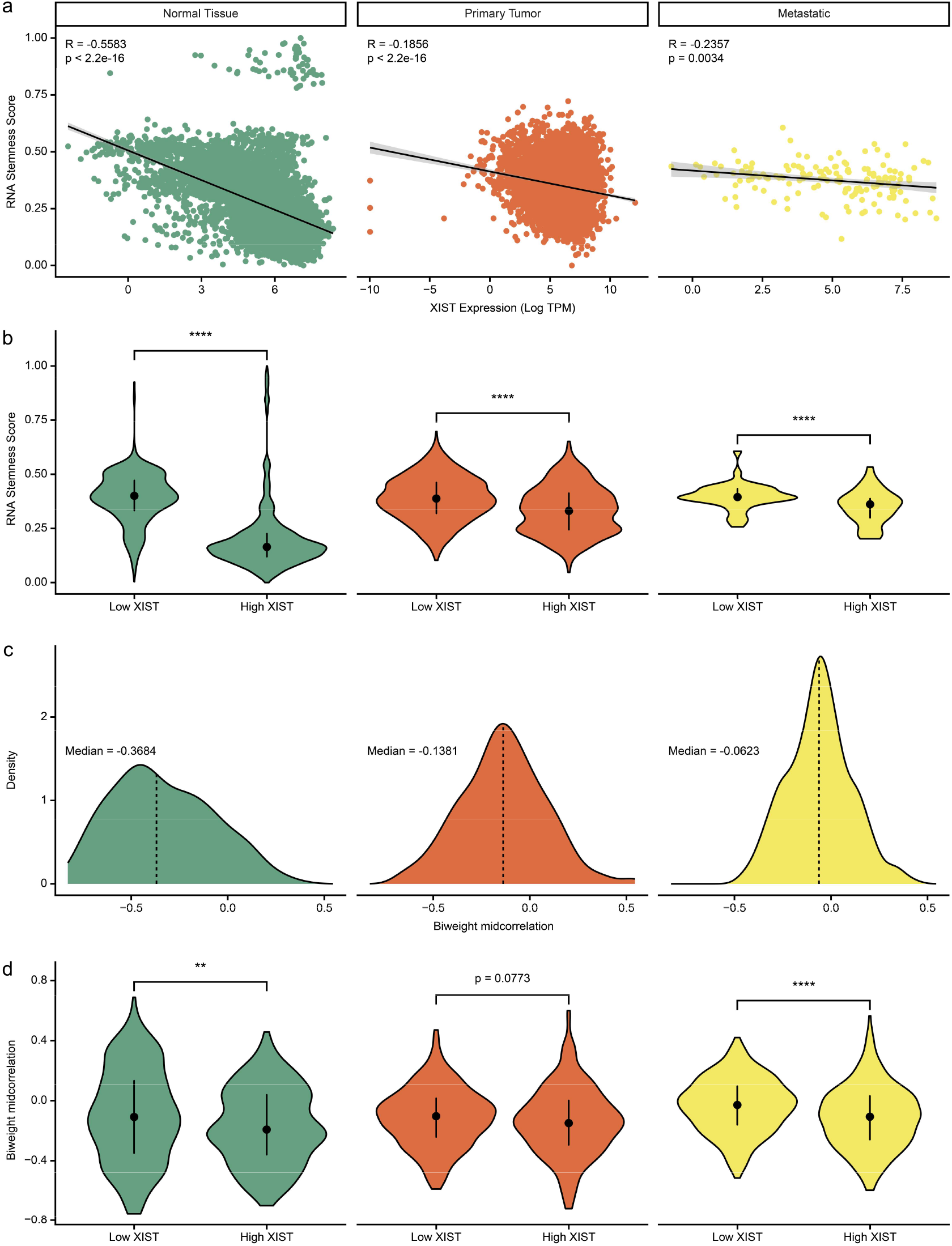
*XIST* and the adhesome show negative correlations with stemness across normal and cancerous states. (a) Scatter plots showing the correlation (calculated by biweight midcorrelation) between *XIST* and stemness score in normal tissue, primary tumor, and metastatic samples. (b) Violin plots comparing stemness scores in low and high *XIST* samples (lower or upper quartile) in normal tissue, primary tumor, and metastatic samples. Data normality was assessed using the Shapiro-Wilk test and statistical analysis was performed by Wilcoxon test. (c) Density plots of adhesome-stemness correlation (calculated by biweight midcorrelation) in normal tissue, primary tumor, and metastatic samples. (d) Violin plots comparing adhesome-stemness correlations in low and high *XIST* samples (lower or upper quartile) in normal tissue, primary tumor, and metastatic samples. Data normality was assessed using the Shapiro-Wilk test and statistical analysis was performed by Wilcoxon test (left panel) and unpaired t test (middle and right panel). **, p < 0.01; ****, p < 0.0001.

To determine whether this relationship extends to the adhesome, we analyzed correlations between stemness and adhesome gene expression across normal, primary tumor and metastatic samples. In all three conditions, the median correlations between adhesome genes and stemness were significantly less than zero (p < 0.001, one-sample Wilcoxon signed-rank test), indicating a predominant negative association (Figure 4c). Given the positive correlation between *XIST* and adhesome gene expression, and the inverse correlations of both *XIST* and the adhesome with stemness, we hypothesized that the anticorrelation between adhesome genes and stemness would be stronger in High-*XIST* samples. Consistent with this prediction, adhesome-stemness correlations were more negative in High-*XIST* samples across normal, primary tumor, and metastatic tissues; this difference was statistically significant in normal and metastatic tissues (Figure 4d). These findings suggest that the *XIST*-associated adhesome network may contribute to suppressing stemness and stabilizing differentiated cellular states, thereby limiting the acquisition of developmental and cancer stem-like properties.

Overall, our analysis demonstrates that both *XIST* and the adhesome help maintain cellular states by suppressing stemness under normal and cancer conditions, highlighting their inter-connected functional roles in tumor progression.

## Discussion

In this study, we identify a previously unrecognized relationship between *XIST* and the adhesome network. Following *XIST* loss, a subset of adhesome genes was significantly differentially expressed; notably, only one of these, *ARHGEF6*, is X-linked, indicating that the majority of affected adhesome genes are autosomal. Although *XIST* is best known for coating the X chromosome and mediating X-chromosome inactivation, its regulatory functions are highly dependent on cellular state. In fully differentiated cells, loss of XIST does not typically lead to widespread reactivation of X-linked genes, and growing evidence indicates that *XIST* can also regulate autosomal transcription (18, 56–58). Accordingly, the deregulation of adhesome-related genes likely reflects both direct and indirect consequences of *XIST* perturbation, acting through a limited set of reactivated X-linked regulators as well as through autosomal targets. This interpretation is consistent with prior reports demonstrating that *XIST* interacts with autosomal loci to modulate their transcription (18, 57, 58), extending its regulatory influence beyond X-chromosome inactivation. The broad adhesome-associated transcriptional changes observed following *XIST* perturbation may therefore arise from multiple mechanisms, including direct *XIST*-chromatin interactions, indirect effects mediated by dysregulated X-linked regulators, and broader alterations in nuclear organization and chromatin compartmentalization.

Our integrative analysis of expression correlations using GTEx and TCGA metadata further emphasizes the broad connection between *XIST* and the adhesome network across normal tissues and primary tumors. We compared *XIST*-adhesome correlations with those of representative lncRNAs, including *MALAT1* and *GAS5*, which have reported functions in modulating adhesion-related cellular processes. For example, *MALAT1* has been shown to promote cell migration, EMT, and metastasis, and *GAS5* has been implicated in regulating vascular smooth muscle cell migration and differentiation (47–49). Interestingly, while these lncRNAs are closely linked to adhesion, *XIST* exhibited the strongest correlations with adhesome genes specifically in normal tissue. This positions *XIST* as a potential contributor to maintaining adhesome homeostasis under physiological conditions, distinguishing it from other lncRNAs. Importantly, we observed that *XIST*-adhesome correlations were reduced in primary tumors, indicating that this regulatory relationship is rewired in malignancy. Additionally, dysregulation of *XIST* has been reported in multiple cancers, where it can influence cell migration, invasion, and EMT (14–16). Together, these findings suggest that both the functional interactions and expression patterns of *XIST* are altered in malignancy, potentially contributing to adhesion-related changes that facilitate tumor progression.

Our findings demonstrate that OVCAR3 cells serve as a biologically relevant model for uncovering *XIST*-associated alterations within the adhesome network. Interestingly, unlike most tissue types in which tumorigenesis typically weakens the *XIST*-adhesome correlation, ovarian tumors maintain a correlation comparable to that of normal ovary, suggesting a tissue-specific association that persists during tumor development. Although ovarian tumors exhibit a relatively modest *XIST*-adhesome correlation compared with other normal and tumor tissues, *XIST* loss in OVCAR3 cells still triggered substantial changes in adhesome gene expression and adhesome-related pathways. This indicates that even a modest correlation is sufficient for *XIST* to play a meaningful regulatory role in modulating the adhesome network. Importantly, because many other normal and tumor tissues show stronger *XIST*-adhesome coupling, the regulatory function of *XIST* may be even more pronounced in contexts where *XIST* maintains tighter coordination with adhesome expression.

Focusing on nuclear-localized adhesome genes enabled us to define an *XIST*-associated adhesome network in which *CTNNB1* functions as a major regulatory hub within the nucleus. Several reports provide mechanistic support for a regulatory connection between *XIST* and the WNT/β-catenin signaling pathway (54, 55). For example, a recent study by Zou et al. demonstrated that *XIST* functions as a competing endogenous RNA for miR-1264 in vascular smooth muscle cells, relieving miR-1264-mediated repression of *WNT5A* and activating WNT/β-catenin signaling, along with increased β-catenin and reduced E-cadherin (55). This *XIST*-microRNA-WNT/β-catenin crosstalk suggests that *XIST* may participate in the adhesome regulatory network by interacting with *CTNNB1* via microRNA. Within this network, adhesome genes such as *CTNNB1* and *ITGB3BP*, as well as *XIST* itself, exhibited lower hazard ratios in tumors with high *XIST* expression. Consistent with this, patients with elevated *XIST* levels displayed improved survival in ovarian cancer (16). Together, these observations suggest that *XIST* is closely associated with adhesome network stability under homeostatic conditions, potentially protecting against aggressive tumor progression. In tumors, altered *XIST* expression may disrupt this *CTNNB1*-centered network, leading to adhesion remodeling, increased motility, and invasion, thereby linking epigenetic regulation, adhesion signaling and patient prognosis.

When investigating the effect of *XIST* on stemness, we found that *XIST* is negatively correlated with stemness across normal tissue, primary tumor, and metastatic samples, indicating that it suppresses stem-like phenotypes in multiple biological contexts. This observation is consistent with published studies showing that *XIST* helps maintain differentiated states in both development and cancer. Deleting *Xist* in female hematopoietic stem and progenitor cells results in defects in lineage-specific differentiation during hematopoiesis, linking *Xist* function to developmental lineage choices (59). Loss of *XIST* impairs mammary stem cell proliferation while promoting tumorigenicity and metastasis (17). Moreover, *XIST* knockdown in OVCAR3 increases stem-like properties and cellular plasticity (16), and downregulation of *XIST* promotes brain metastasis via activation of EMT and stemness (60). Importantly, our work extends these findings by showing that the suppressive role of *XIST* in stemness is not limited to isolated experimental systems but is broadly supported across large tissue cohorts (GTEx and TCGA datasets).

Beyond *XIST*, several core adhesome genes have been implicated in suppressing stemness across multiple studies. For example, integrins are essential for maintaining epithelial differentiation and tissue integrity, and loss of integrin function has been shown to promote dedifferentiation and the acquisition of stem-like properties in mammary epithelial cells (61). Similarly, Focal adhesion kinase (*FAK/PTK2*) and Rho-ROCK signaling – two central adhesome hubs – are critical regulators of multilineage differentiation in human adipose stem cells (62). E-cadherin, a central component of cell-cell adhesion, also constrains cellular plasticity; its loss drives EMT and stemness across multiple cancer types (63). Together, these experimental observations support our broad finding that adhesome genes are generally anti-correlated with stemness in both GTEx and TCGA analyses. Importantly, because *XIST* expression is strongly correlated with adhesome genes in normal but is markedly weakened in tumors, this disruption may decouple epigenetic regulation from adhesion signaling, thereby promoting plasticity, stemness acquisition, and invasion potential in tumor cells. This context-dependent regulation indicates that *XIST*’s role in the adhesome network is not static but dynamically altered during tumorigenesis.

A limitation of this study is that the analyses are primarily associative and therefore do not establish causal relationships between *XIST* and specific adhesome genes. Although we observed strong correlations across large datasets and consistent transcriptional changes following *XIST* knockdown, these findings alone cannot determine whether *XIST* directly regulates adhesome genes or stemness-related pathways. Additional functional studies, such as targeted perturbation and rescue experiments, will be required to distinguish direct regulatory mechanisms from indirect effects. Another limitation is that several analyses aggregate data across diverse tissue and tumor types from GTEx and TCGA. While this approach increases statistical power and enables the identification of broad, conserved patterns, it may obscure tissue-specific regulatory relationships and influence the magnitude of observed correlations. Future studies incorporating tissue- and cancer-type-specific analyses will therefore be necessary to refine these associations and elucidate underlying mechanisms.

Overall, our findings establish *XIST* as a central component of the adhesome network, with its dysregulation linked to tumor progression. This study uncovers a previously unrecognized epigenetic-adhesion regulatory axis that becomes remodeled in tumors, influencing cell adhesion, cellular plasticity, and cancer progression.

## Materials and Methods

### Differential Gene Expression and Gene Ontology Analysis

OVCAR3 *XIST* knockdown RNA-seq profiles have been previously reported and deposited in the Gene Expression Omnibus (GEO: GSE271117) (16). As previously described, two independent knockdown clones, X7 (sgXIST7) and X9 (sgXIST9) were generated using distinct guide RNAs targeting different regions of the *XIST* promoter (sgXIST7: 5’-GCAGCGCTTTAAGAACTGAA-3’; sgXIST9: 5’-GCCATATTTCTTACTCTCTCG-3’), with the control clone generated in parallel using an sgRNA targeting the Gal4 promoter (sgCtl: 5’-GAACGACTAGTTAGGCGTGT-3’). For the present analysis, normoxic samples were used, including the control (n = 3) and two independent *XIST*-knockdown clones (X7 and X9; n = 3 each). Differential expression analysis was based on the DESeq2 output from the prior study Dataset S01(16). In contrast to the prior analysis, which used a strict cutoff of |log_2__FoldChange| > 1 (corresponding to a 2-fold change), the present analysis defined differentially expressed genes (DEGs) using a more permissive threshold of |log2_FoldChange| > 0.58 (corresponding to a 1.5-fold change) and adjusted p value < 0.05. The 1.5-fold cutoff was selected to increase sensitivity for detecting biologically meaningful *XIST*-dependent transcriptional changes and to capture subtle but relevant alterations in the regulatory landscape that may be excluded by a more conservative 2-fold threshold. Thus, this cutoff balances biological relevance with statistical stringency(64, 65). The DEG list and adhesome gene list are provided in Supplementary Tables 1 and 2 (20), Differentially expressed adhesome genes were identified by overlapping these lists. To minimize potential sgRNA-specific off-target effects, we further restricted analyses to adhesome genes that were consistently differentially expressed in both knockdown clones. This cross-validation strategy aligns with established CRISPR experimental design principles, in which concordant effects across distinct guides increase confidence that observed transcriptional changes reflect the intended perturbation rather than guide-specific artifacts (66).

Gene ontology (GO) enrichment analysis for Gene Ontology Biological Process (GOBP) terms was performed using the gprofiler2 R package. In contrast to the prior analysis, which used all annotated human genes as the background, the present DEG lists were tested against a custom background gene set consisting of expressed genes, defined as genes with a total raw count greater than 50 across all samples. Enrichment p values were calculated using a hypergeometric test with multiple testing correction by the false discovery rate (FDR) method, and significant GO terms were defined as those with FDR-adjusted p-value < 0.05.

### Correlation Analysis

A panel of well-characterized regulatory lncRNAs with diverse reported functions was selected to provide a biologically relevant reference framework for evaluating relative correlation strengths with adhesome gene expression. Within this lncRNA panel, *GAS5* and *MALAT1* have been reported to be involved in adhesome-related pathways (47–49). Gene expression profiles for lncRNAs and adhesome genes were obtained from GTEx and TCGA metadata (TCGA data were extracted from UCSC’s Xena Data Portal). Correlations between lncRNAs and adhesome genes were calculated using biweight midcorrelation in R, a median-based statistical correlation method that is less sensitive to outliers and therefore well suited for gene expression comparisons (67).

To evaluate whether the observed correlation between *XIST* and the adhesome gene set exceeded random expectation, a permutation-based approach was applied. Specifically, 2,000 random gene sets of equal size (n = 334) were sampled from all genes (excluding *XIST*), and their median correlations with *XIST* were calculated using the same method. As an additional control, randomly sampled X-linked gene sets of equal size (excluding *XIST*) were analyzed. The observed median correlation was then compared with the distribution of median correlations derived from these random gene sets.

### *XIST*-Associated Adhesome Network Construction

The *XIST*-associated adhesome network was constructed using the STRING (Search Tool for Retrieval of Interacting Genes/Proteins) database (68) and visualized with Cytoscape (69). Network nodes represent adhesome genes potentially localized in the nucleus (annotated by subcellulaRVis (53)) and ranked within the top 10% by correlation with *XIST* expression.

### Hazard Ratio Analysis

Patients were stratified into high and low *XIST* expression groups based on quantile expression. Univariate Cox proportional hazards regression was performed to estimate hazard ratios for *XIST* and each adhesome gene, and statistical significance was assessed using the Wald test. To ensure sufficient statistical power and model stability, all TCGA cancer types were pooled for the Cox proportional hazards analysis, as individual cancer types did not provide enough samples for reliable model convergence.

### Stemness Score Analysis

RNA stemness scores were defined using the stemness index framework described by T. M. Malta et al., which applies a machine-learning-based model trained on stem cell gene expression profiles to quantify the degree of oncogenic dedifferentiation in each sample (70). Stemness scores for female normal tissue were calculated by applying the TCGAanalyze_Stemness function (70) to GTEx expression data. Stemness scores for female primary tumor and metastatic samples were obtained directly from TCGA metadata, where they had been previously calculated using the same framework. Correlations between stemness scores and *XIST* or adhesome gene expression were assessed using biweight midcorrelation in R.

## Code Availability

All R scripts used in this analysis were deposited on GitHub (https://github.com/downinglab/chen-origer-xist-adhesome).

## Acknowledgements

This work was supported by National Institutes of Health NIH R01GM141424 (S.S.) and by the National Institute of Neurological Disorders and Stroke of the NIH T32NS082174 (N.O.). The content is solely the responsibility of the authors and does not necessarily represent the official views of the National Institutes of Health. Research was supported in part by the National Cancer Institute of the National Institutes of Health under award number P30CA062203 (T.L.D.) and the UC Irvine Comprehensive Cancer Center using UCI Anti-Cancer Challenge funds (S.S., T.L.D.).

