## Supplementary Figures for "*XIST* Is a Key Modulator Associated with the Adhesome Network"

### Supplemental Figure 1

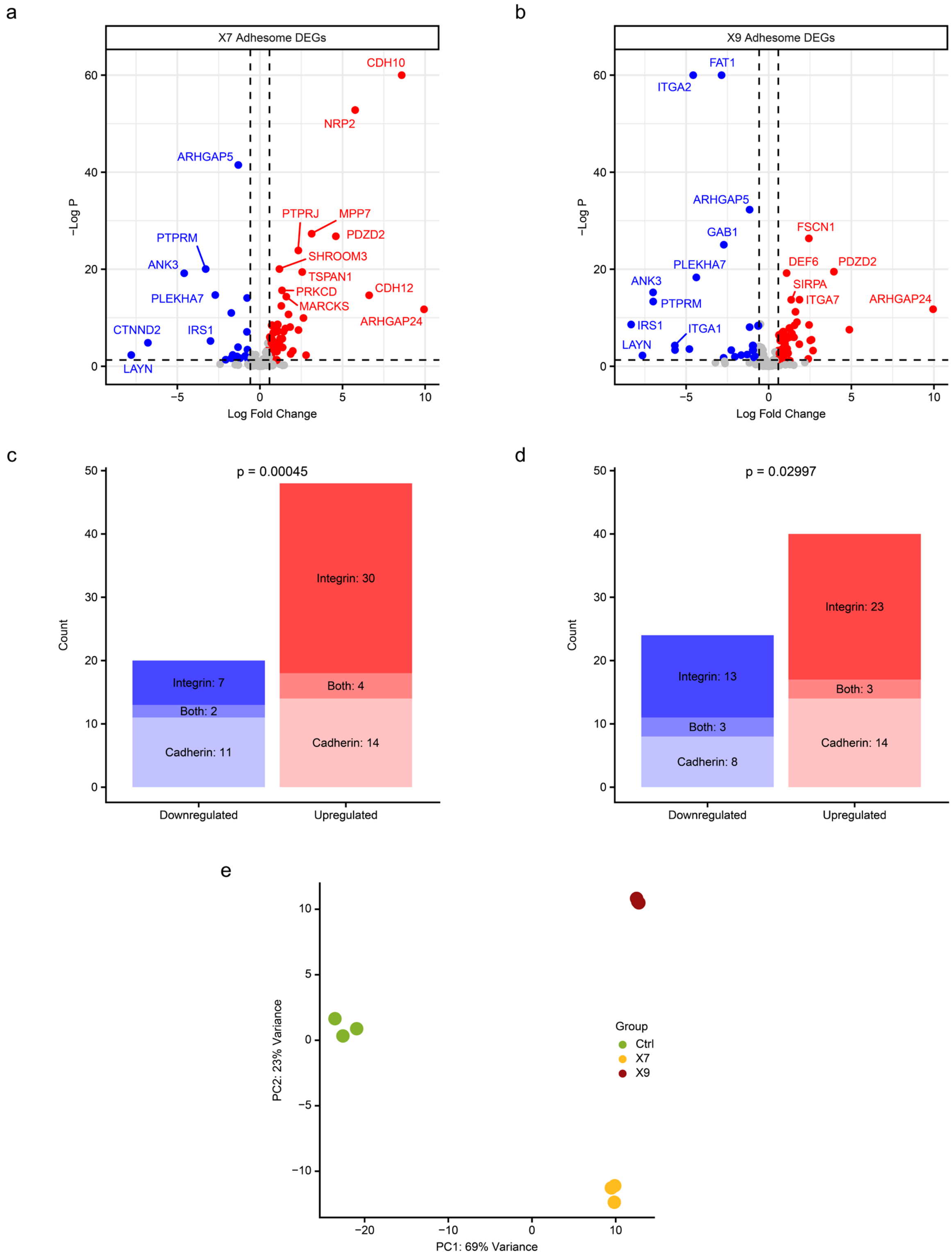

**Figure S1.** (a-b) Volcano plots showing the differentially expressed adhesome genes after *XIST* knockdown in OVCAR3 clones X7 (a) and X9 (b). Adhesome genes with |log2FoldChange| > 3 or -log10(padj) > 20 were labeled with their gene names. (c-d) Counts of downregulated and upregulated adhesome genes classified as integrin-associated (Integrin), cadherin-associated (Cadherin), or shared between both categories (Both) in *XIST*-knockdown OVCAR3 clones X7 (c) and X9 (d); statistical comparisons were performed using a one-tailed binomial test. (e) Principal component analysis (PCA) of all nine samples, including control cells (Ctrl, n = 3) and two *XIST* knockdown clones (X7 and X9, n = 3 each), performed using DESeq2.

### Supplemental Figure 2

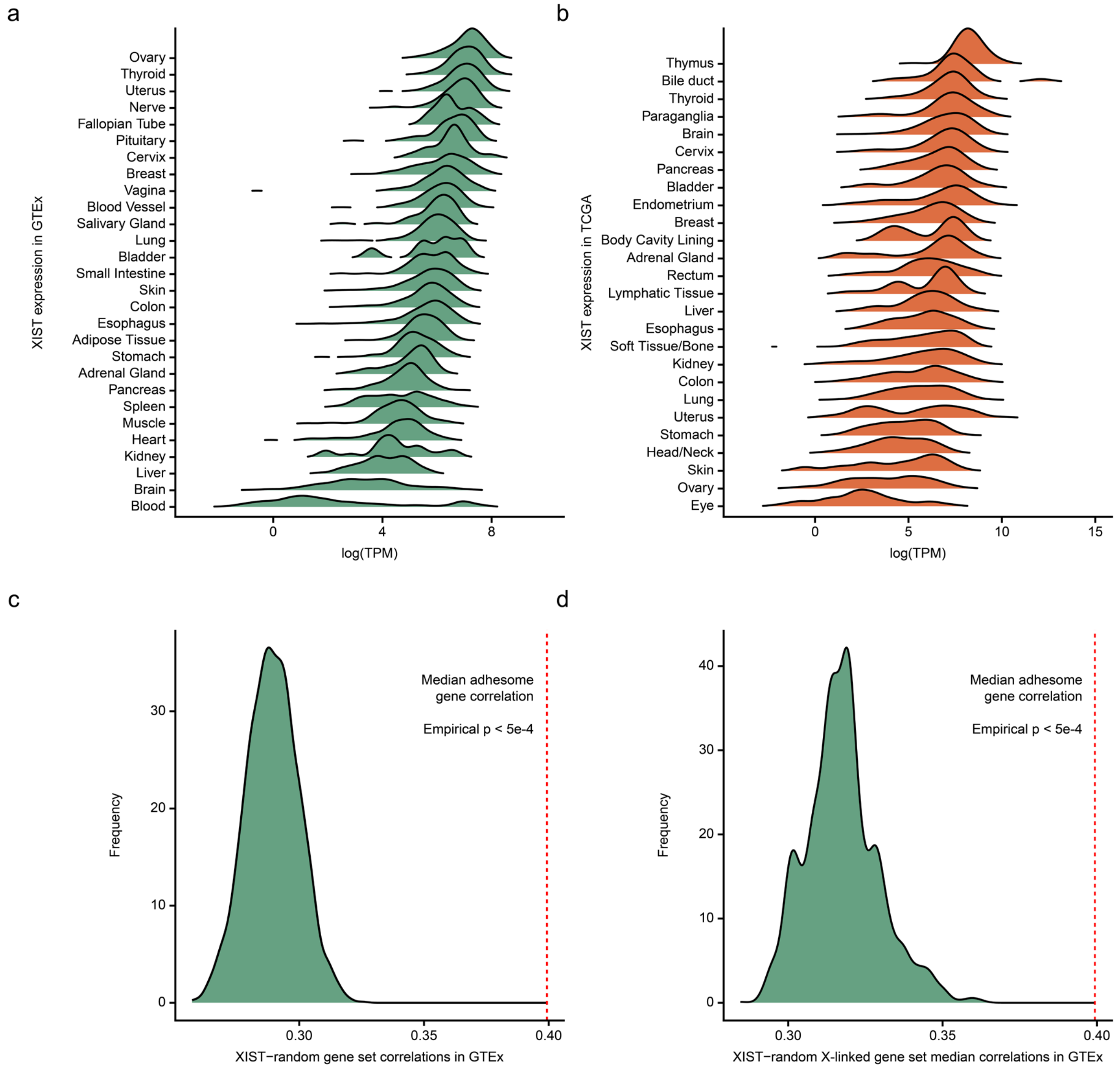

**Figure S2.** (a-b) *Xist* expression levels ($log_2$(TPM+0.001)) across tissue types in GTEx (a) and TCGA (b). (c-d) Distribution of median correlations between *XIST* and 2,000 randomly generated gene sets (c) or X-linked gene sets (d) matched in size to the adhesome gene list (n = 334). The observed median correlation between *XIST* and adhesome (R = 0.3993; red dashed line) exceeds that of all random permutations. The empirical p-value was calculated as the fraction of random gene sets whose median correlation values were greater than or equal to the observed median adhesome gene correlation, using the formula (1 + k) / (1 + N), where k is the number of permutations ≥ the observed value and N is the total number of permutations.

Supplemental Figure 3

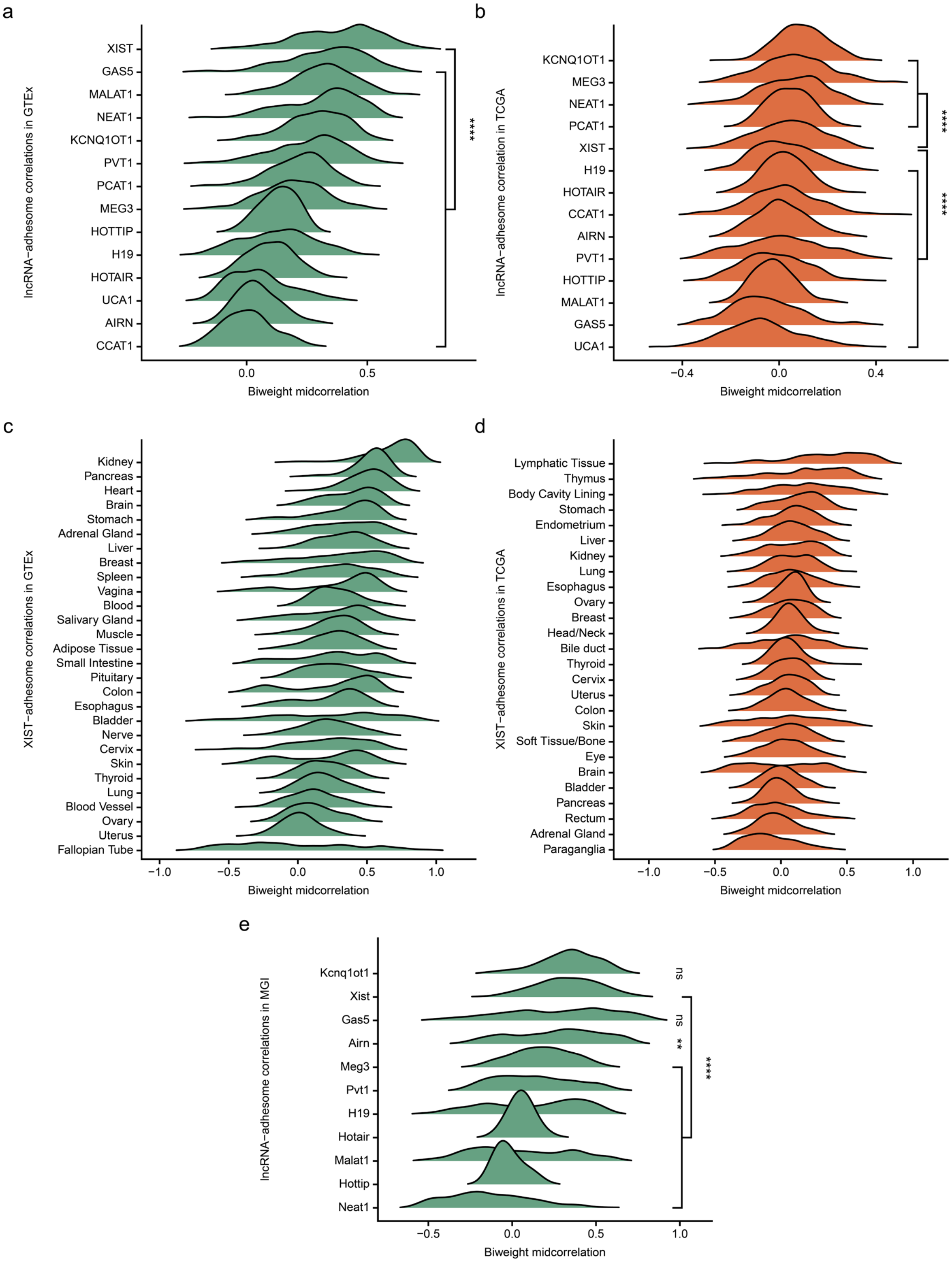


**Figure S3.** (a-b) Correlations between lncRNAs and adhesome genes in female normal tissues (a) and female primary tumors (b); statistical significance was assessed using a one-tailed Wilcoxon test comparing *Xist* with each of the other lncRNAs. (c-d) Correlation between *XIST* and adhesome genes across tissue types in female normal tissues (c) and female primary tumors (d). (e) Correlations between lncRNAs and adhesome genes in mouse normal tissues; statistical significance was assessed using a one-tailed Wilcoxon test comparing *Xist* with each of the other lncRNAs. ns, not significant; **, $p < 0.01$; ****, $p < 0.0001$.
